# Precision measurement of non-conscious avoidance reactions using 3D tracking: Validation across olfaction and vision

**DOI:** 10.64898/2026.04.30.721883

**Authors:** Evelina Thunell, Elisa Dal Bò, Frans Nordén, Artin Arshamian, Marie Michael, Supreet Saluja, Hedvig Kjellström, Arnaud Tognetti, Johan N. Lundström

**Affiliations:** Department of Clinical Neuroscience, Karolinska Institutet, Nobels väg 9, 171 77 Stockholm, Sweden; Department of General Psychology, University of Padua, Via Venezia 8, 35131 Padua, Italy; Department of Comparative Cultural Psychology, Max Planck Institute for Evolutionary Anthropology,Deutscher Platz 6, 04103 Leipzig, Germany; Max Planck School of Cognition, Max Planck Institute for Human Cognitive and Brain Sciences, Stephanstrasse 1a, 04103 Leipzig, Germany; Charles Perkins Centre, The University of Sydney, Sydney, NSW 2050, Australia; Department of Robotics, Perception and Learning, KTH Royal Institute of Technology, 100 44 Stockholm, Sweden; CEE-M, University of Montpellier, CNRS, INRAE, Institut Agro, Montpellier, France; Monell Chemical Senses Center, 3500 Market Street, PA Philadelphia, USA

**Keywords:** approach-avoidance behavior, postural reactions, olfactory perception, visual perception

## Abstract

One of our sensory systems’ key functions is to detect threats in the environment. Sensory information eliciting negative emotions, such as fear or disgust, triggers instinctive avoidance reactions. This core survival mechanism is believed to be expressed as subtle non-conscious postural reactions, even when participants are instructed to stand still. Such avoidance behavior has mainly been studied using indirect measures that make participants aware of their posture (e.g. force-plate based methods) or measures that depend on explicit cognitive tasks, like moving a joystick to indicate an urge to approach or avoid the stimulus; experimental tasks with limited ecological validity and generalizability. Therefore, despite the importance of this basic survival strategy, its underlying mechanisms are still poorly understood. Here, we used a novel 3D-camera-based method allowing direct but implicit measures of postural reactions with high precision. Participants are aware that they are being filmed but, crucially, are not informed that distance measures are obtained. We assessed this ecologically valid measure of approach/avoidance responses in two different sensory modalities: olfaction and vision. Participants were standing upright while exposed to either olfactory or visual stimuli and verbally rating their perceived valence in each trial. In response to subjectively unpleasant odors and images, participants moved away from the stimulus source, as compared to pleasant stimuli. These results demonstrate a putative modality-independent early proxy for avoidance behavior in response to perceived negative valence. Considering its face validity and general applicability, this novel experimental method presents new possibilities for assessing non-conscious approach-avoidance responses in humans.

## Introduction

The ability to evaluate sensory information as either beneficial or harmful is a fundamental evolutionary function that promotes survival by guiding approach and avoidance behaviors (McNaughton et al., 2016). This core evaluative process is ubiquitous across phyla, from the simple phototaxis of single-celled organisms to the complex decision-making of humans (Arshamian et al., 2017; Elliot et al., 2013). In humans, sensory information associated with potential threats, such as the sight of an aggressor (Monéger et al., 2025) or the smell of blood (Arshamian et al., 2017), often gives rise to powerful negative emotions like fear or disgust, which serve as potent avoidance cues. Many theories of emotion suggest a fundamental connection between emotions and action tendencies (Frijda, 1986, 2009) for mobilizing the organism to respond adaptively to emotionally significant stimuli in the environment (Lang et al., 1990). The biphasic theory of emotion provides a robust framework for this link, positing that human motivation is organized around two primary systems: an appetitive system that facilitates approach toward pleasant or life-sustaining stimuli, and a defensive system that triggers withdrawal from unpleasant or threatening stimuli (Lang, 1995; Lang et al., 1990). These motivational systems are thought to be evolutionarily conserved and are reflected in adaptive physiological and behavioral responses to a dynamic environment (Arshamian et al., 2017; Elliot, 2006; Fricke & Vogel, 2020; Koshland, 2002; Rutherford & Lindell, 2011) (Bradley et al., 2001; Harrison et al., 2015). The associated behaviors are expressed both over long timescales and critically, over short timescales, as subtle non-conscious postural approach-avoidance tendencies. However, there has so far been a lack of methods that can accurately measure the latter effects with high face validity.

Despite the clear ecological value of the basic survival strategy of approach and avoidance behavior, its underlying mechanisms in humans have been surprisingly difficult to study (Monéger et al., 2025). Much of our knowledge comes from animal research, where approach and avoidance behaviors are more readily observable (Kirlic et al., 2017). Research in humans has yielded mixed and often ambiguous findings, largely due to methodological challenges (Monéger et al., 2025). The research field has relied on indirect measures where participants are explicitly prompted for an approach-avoidance reaction (e.g., joystick or button-press tasks) and using tasks that have low ecological validity (Monéger et al., 2025). So far, the most ecologically valid method has been to measure postural sway using force plates; however, this method risks making the participant aware that their posture or balance is being monitored, and the empirical evidence has been inconsistent with substantial heterogeneity, as revealed in a recent meta-analysis (Monéger et al., 2025). Thus, available methods risk measuring conscious, cognitive strategies, demand characteristics, or deliberate actions rather than the automatic motor adjustments that are the presumed approach-avoidance output of the core motivational systems (Chen & Bargh, 1999; Roelofs, 2017).

To overcome these limitations and capture automatic non-conscious approach-avoidance responses in a direct and ecologically valid way, we employed a novel experimental setup based on a 3D camera measuring subtle, frame-to-frame displacements of the participant toward or away from the target, with high precision. Participants were standing upright while we recorded their approach-avoidance reactions (distance between the 3D camera and their nose tip) to visual or olfactory stimulation. Recognizing that the emotional response to a stimulus is highly subjective and can even vary across trials, we probed the perceived valence on a trial-by-trial basis to ensure that our measure was tied to the participants’ true experience.

## Methods

### Participants

For both experiments, we recruited participants reporting that they had normal or corrected-to-normal vision, not used facial botox in the past year (to allow for future analyses of facial expressions), and no balance problems, ADHD, or neurodegenerative disorders. In the olfactory experiment, we added the criteria to have a self-assessed normal sense of smell and no nasal congestion on the day of testing, and instructed participants not to eat or drink anything else than water for the 30 minutes leading up to the experiment.

All procedures were in accordance with the Declaration of Helsinki and approved by the Swedish Ethical Review Authority (Dnr 2021-06766), and all participants provided written informed consent prior to enrolling in the experiment. The visual experiment was preregistered at https://aspredicted.org/tsxn-rryw.pdf

#### Visual experiment

For the visual experiment, we recruited 98 participants via an online recruitment system. We excluded the data from 22 of these due to less than half of the trials being usable (we excluded trials where the face detection algorithm failed, and with missing or inaudible verbal ratings). The data from another 5 participants were excluded due to excessive swaying, changing the weight back and forth between the feet, head and face twitching, shoulder movement, yawning, and stretching, as detected post-hoc in the recorded videos. These behaviors were seen mainly in the 5 excluded participants with rare occurrences for other participants. The final sample thus consisted of 76 participants (47 women, 29 men), aged 18-50 (M = 28.8 years, SD = 7.1). Methods and analyses were in accordance with our pre-registration, unless otherwise explicitly stated.

#### Olfactory experiment

Seventy individuals participated in the olfactory experiment, recruited the same way as for the visual experiment. To avoid including participants with functional anosmia, we assessed their olfactory ability using five common odors from the standard Sniffin’ Sticks (Sniffin’ Sticks, Burghart, Germany) identification test (lemon, licorice, cinnamon, garlic, and banana, presented in felt pens). Participants performed a 4-alternative forced-choice cued identification task, and we included participants who correctly identified at least three out of the five odors. One participant failed this olfactory test criterion and was thus not included. We used the same criteria for data exclusion as in the Visual experiment: The data from 22 participants were excluded due to less than half of the experimental trials being usable (because of the face detection algorithm failing or missing/inaudible verbal ratings); no participants displayed excessive movements. The final sample thus consisted of 47 participants (29 women, 18 men). The participants were between 19 and 38 years old (M = 26.7, SD = 4.3).

### Stimuli and task

#### Visual experiment

The stimuli consisted of 108 images from the International Affective Picture System (IAPS; (Lang et al., 2005) chosen to elicit pleasant, neutral, and unpleasant percepts, based on the standardized ratings of the database. Thirty-six pleasant images were chosen based on their normative valence ratings (M = 7.27, SD = 1.55 on a scale 1-9), as well as 36 neutral images (M = 5.15, SD = 1.20) and 36 unpleasant images (M = 2.70, SD = 1.61). We purposefully did not include the most negatively rated images, to avoid very negative reactions or freeze-like responses from the participants. The IAPS codes for the images can be found in the Supplementary Material. The images were square in shape with a side length of 27 cm presented at around 60-75 cm distance on a screen.

Each image was presented for 2000 ms, after which participants were prompted to rate their perceived valence of the stimulus by speaking it out loud. To prompt this task, we displayed a 9-point scale for 4000 ms (from 1 - very unpleasant, to 9 - very pleasant). After the offset of the rating scale, there was a random interval between 1 and 3 seconds before presentation of the next image. Each image was presented to each participant once in a random order. To avoid fatigue, the experiment was divided into three 6-minute blocks, each with 12 images from each of the three categories pleasant, neutral, and unpleasant. Before initiating the experiment, participants practiced the task by rating three images that were not included in the experimental image set.

#### Olfactory experiment

We used a computer-controlled olfactometer (Lundström et al., 2010) to present 6 different odor qualities to participants: rotten food (diethyl disulfide), urine (complex mixture), parmesan/vomit (butyric acid), pineapple (ethyl butyrate), grass (cis-3-hexenol), and lilac (complex mixture). The odors were selected to represent a large range of the valence scale for most participants. Each odor was presented 16 times, resulting in 96 odor presentations in total. These were divided into 4 blocks lasting around 7 minutes each (24 trials per block in a randomized order). To prevent participants from predicting the onset of an odor stimulus while ensuring a clear percept, the odor was triggered (without participants’ knowledge) based on their own breathing cycle. Each stimulus was presented for 2000 ms, together with a fixation cross presented on the screen at about 60-75 cm distance which remained for 1000 ms after stimulus offset. After another 500 ms of blank (background gray) screen, the 9-point valence rating scale was presented for 4000 ms (from 1 - very unpleasant, to 9 - very pleasant) and, after 500 ms of blank screen, a scale to rate the intensity of the odor stimulation (from 1 - very weak, to 9 - very strong). Half of the participants were instead prompted to rate odor intensity first and thereafter odor valence. Last, a blank screen was presented for 500 ms followed by the fixation cross for a random duration of between 1000 and 2000 ms. After this, the next odor was released at the lowest point of the inhalation phase in the subsequent breath, to ensure that the odors were likely presented during inhalation. Participants’ breathing cycles were measured with a nasal cannula connected to a Powerlab unit via a Spirometer Pod (Powerlab 16/35, ADInstruments). The respiration signal used to trigger odor delivery was analyzed online using LabChart recording software (ADInstruments). The odors were embedded in an ongoing airstream 10 cm in front of the nose. We used a mini photoionization detector (PID; Aurora Scientific Inc, model 200085) to measure the onset time, i.e. travel time from odor nozzle to the nose, of each of the six odors. As stimulus onset time for each odor, we used the average delay of 50% intensity of the plateau level (rotten food: 333 ms, urine: 398 ms, parmesan/vomit: 430 ms, pineapple: 623 ms, grass: 456 ms, lilac: 490 ms). These values were based on 10 measurements per odor done in the same conditions as during our experiment. Before launching the experiment, participants practiced the task by rating three trials of clean air.

### Procedure

Participants were not informed that approach–avoidance behavior was being assessed, either before or during the experiment. They were instructed to stand naturally with their arms at their sides on a marked line on the floor (see Figure 1), and to avoid unnecessary movements. High-resolution video and participants’ verbal ratings were recorded using a Panasonic video camera. The distance to the nose of the participant was measured with a 3D camera (Intel RealSense D435 or Intel RealSense D455) placed directly above the screen. The 3D camera recorded at a sampling frequency of 60 Hz and was tuned for maximum accuracy at 60 cm distance, resulting in an RMS depth error less than 1 mm (Grunnet-Jepsen et al., 2019). This method replaced the Apple iPhone camera-based method mentioned in the pre-registration of the Visual experiment, as it has better resolution and usability. The height of both the screen and cameras was adjusted to each participant’s height.

**Figure 1.**
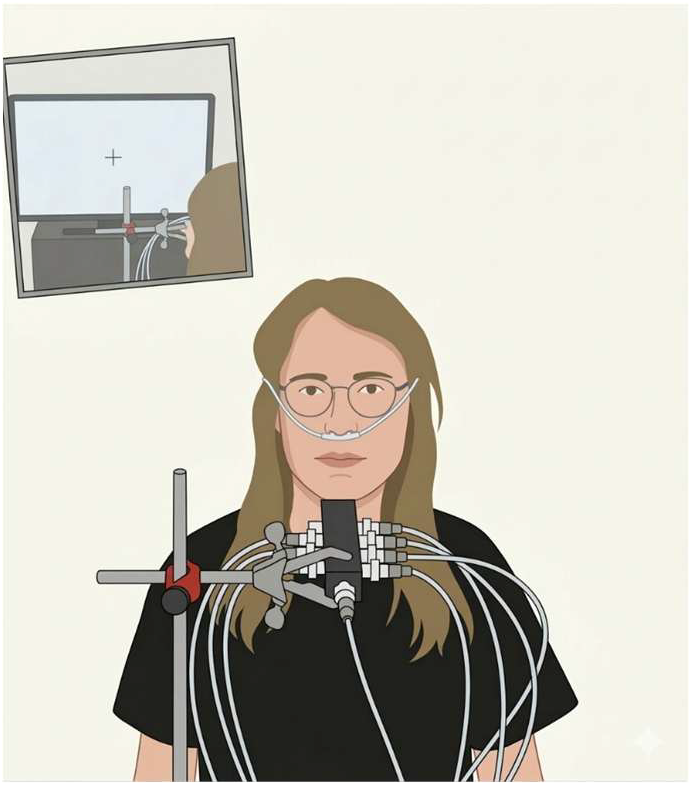
Schematic of the experimental setup in the olfactory experiment; view from the camera. Participants were standing upright in front of the screen and 3D camera and the odor stimulation was administered by an olfactometer. In the visual experiment, the pictures were presented on the screen.

For temporal alignment between stimulus presentation and recorded data, a mirror mounted on the wall behind the participant reflected part of the screen, which was captured by the video recordings. In between blocks, participants were invited to sit down to avoid exhaustion. Last,participants filled out surveys that were not analyzed in this study (the big five inventory, a health and sleep questionnaire, the state-trait anxiety inventory, and the patient depression questionnaire PHQ-9).

### Analysis

Distance data recorded by the 3D camera were pre-processed by interpolating outlier trials (missing values and values >0.5 m from the trial median value) and down-sampling to 10Hz. To assess postural approach/avoidance reactions to the stimulation, we extracted epochs spanning from 200 ms before stimulus onset, until 4000 ms after stimulus onset. The 200 ms period before stimulus onset was used for baseline correction. Stimulus onset (T = 0 in the epoch) was defined as image onset in the Visual experiment, and as 50% of the maximum odor plateau value in the Olfactory experiment to account for varying delivery times across odors.

To account for individual variations in rating-scale usage, and variations in valence perception across individuals and trials, emotional categories were defined separately for each participant based on their trial-to-trial ratings, rather than on predefined stimulus categories. In the visual experiment, where each image was presented once, category assignment therefore depended on each participant’s subjective valence rating, meaning that the same image could fall into different categories across participants. In the olfactory experiment, this approach accounted not only for individual differences in valence ratings across the six odors, but also across the 16 presentations of each odor. Trial-by-trial verbal ratings were coded offline and used to classify trials into Pleasant, Unpleasant, or Neutral conditions for each participant. To that aim, an adaptive procedure was applied such that approximately one third of each participant’s trials were assigned to each category. Trials rated as 1 were classified as Unpleasant and trials rated 9 as Pleasant. Additional trials were then included by iteratively expanding the rating range toward the center of the scale (Unpleasant: 2,3, then 4; Pleasant: 8,7, then 6), provided that including the next rating level did not cause the category to exceed the target proportion. If either Pleasant or Unpleasant remained empty after this procedure, trials rated as 5 were assigned to the empty category. All remaining trials were classified as Neutral. As a result, ratings ≤4 were never classified as Pleasant and ratings ≥6 were never classified as Unpleasant. In the Olfactory experiment, this procedure was conducted after excluding trials in which participants rated odor intensity as the minimum value 1 (54 trials in total, M = 1.15, SD = 2.24).

We used a cluster-based permutation test to assess differences between the Unpleasant and Pleasant conditions across the epoch, excluding the pre-stimulus baseline period (2-sided dependent test with 10000 permutations), as implemented in the Matlab function permutest.m. This method controls for multiple comparisons across time, while considering the smoothness of the signal. In our preregistration, we also mentioned exploratory tests contrasting Pleasant and Unpleasant images against zero. However, we instead added exploratory tests contrasting Pleasant and Unpleasant images against Neutral.

For comparison, we also grouped trials by the pre-defined categories. These were defined by the normative rating data that we used to select the images in the Visual experiment (Unpleasant, Neutral, and Pleasant), and based on odor identity in the Olfactory experiment (the 6 different odors).

## Results

As predicted, participants leaned away from subjectively unpleasant stimuli as compared to subjectively pleasant stimuli, and this pattern was observed in both modalities (vision and olfaction).

In the Visual experiment, we found a postural avoidance reaction to subjectively Unpleasant images as compared to subjectively Pleasant images (preregistered test) from 1500 ms until the end of the epoch, i.e. 4000 ms after stimulus onset (Fig. 2A; cluster-based permutation testing). Participants’ ratings were for the conditions Unpleasant: 1.9 ± 0.6 (M ± SD), Neutral: 5.0 ± 0.4, and Pleasant: 7.4 ± 0.6. When contrasting Unpleasant against Neutral (exploratory test), the significant times were 2800 ms and 3000-3500 ms after stimulus onset, and for the contrast between Pleasant and Neutral (exploratory test) the significant times were 2100-2600 ms and 2900 ms after stimulus onset.

**Figure 2.**
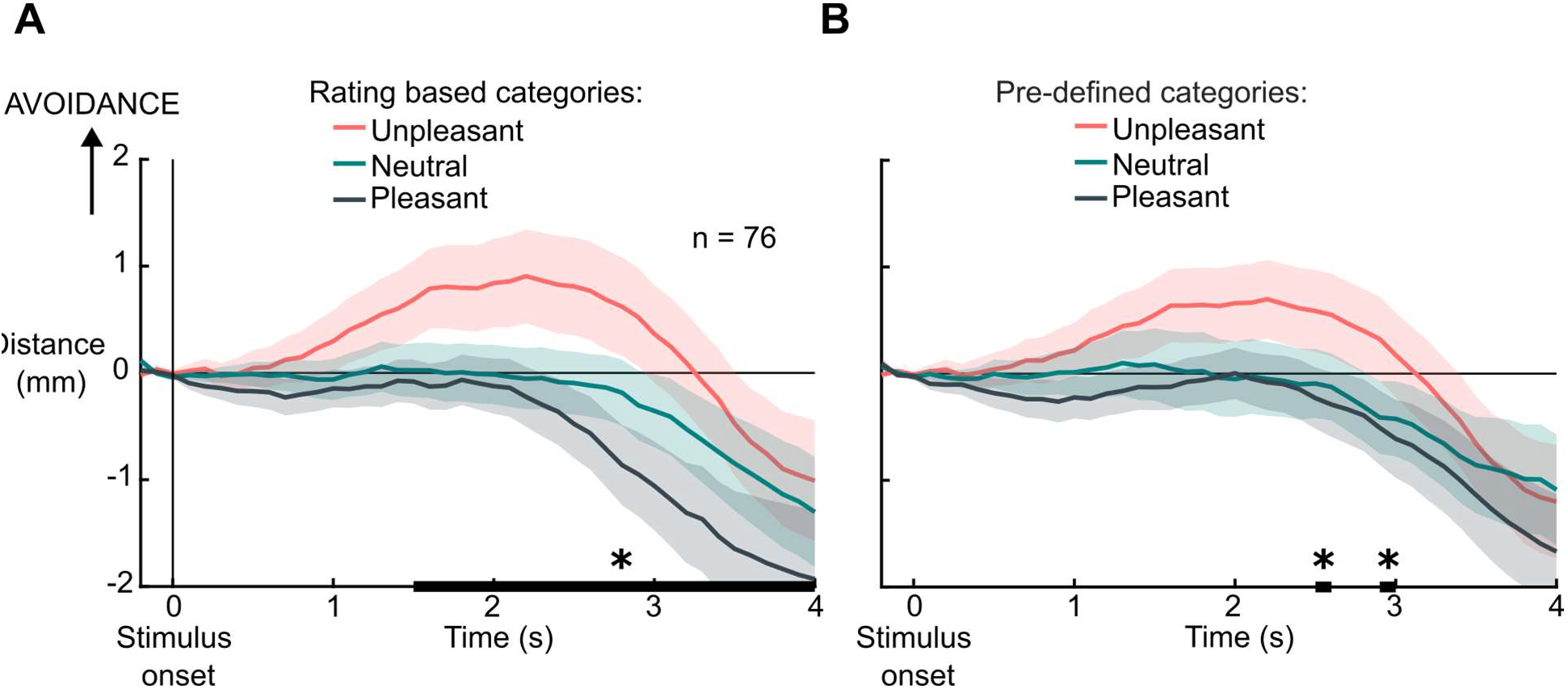
Postural reactions to images. A) Postural reactions to Unpleasant, Neutral, and Pleasant stimulation as defined by participants’ individual trial-by-trial valence ratings. The black thick lines mark clusters of statistically significant time points of difference between Unpleasant and Pleasant trials.Stimulus duration was 2 s. Error bars depict ±1 SEM. B) For comparison, the postural reactions are grouped by the pre-defined (normative) categories Unpleasant, Neutral, and Pleasant.

For comparison, we also analyzed the data grouped by the normative valence ratings from the IAPS database. For these pre-defined categories, there were only two short timepoints of significant difference between Unpleasant and Pleasant, at 2500 ms and 2900 ms after stimulus onset (Fig 2B; cluster-based permutation testing). Our participants’ ratings were for the trials of the predefined categories Unpleasant: 2.3 ± 0.7 (M ± SD), Neutral: 5.2 ± 0.4, and for Pleasant: 6.7 ± 0.7.

In the Olfactory experiment, we found postural avoidance to subjectively Unpleasant odors as compared to Pleasant odors at 300 ms, and 600-4000 ms after stimulus onset (Fig. 3A; cluster-based permutation testing). When instead compared to Neutral, Unpleasant odors elicited an avoidance response in a slightly shorter interval: 1100-4000 ms after stimulus onset. A statistically significant albeit minor approach response to Pleasant odors as compared to Neutral odors was seen at 0-300 ms and 600 ms post stimulus onset. Participants’ valence ratings were for the categories Unpleasant: 2.33 ± 0.66 (M ± SD), Neutral: 4.66 ± 0.82, and Pleasant: 7.16 ±0.73. The intensity ratings did not show the same pattern but were instead comparable between categories: Unpleasant 6.21 ± 1.49, Neutral: 5.21 ± 1.20, and Pleasant: 6.02 ± 1.14.

**Figure 3.**
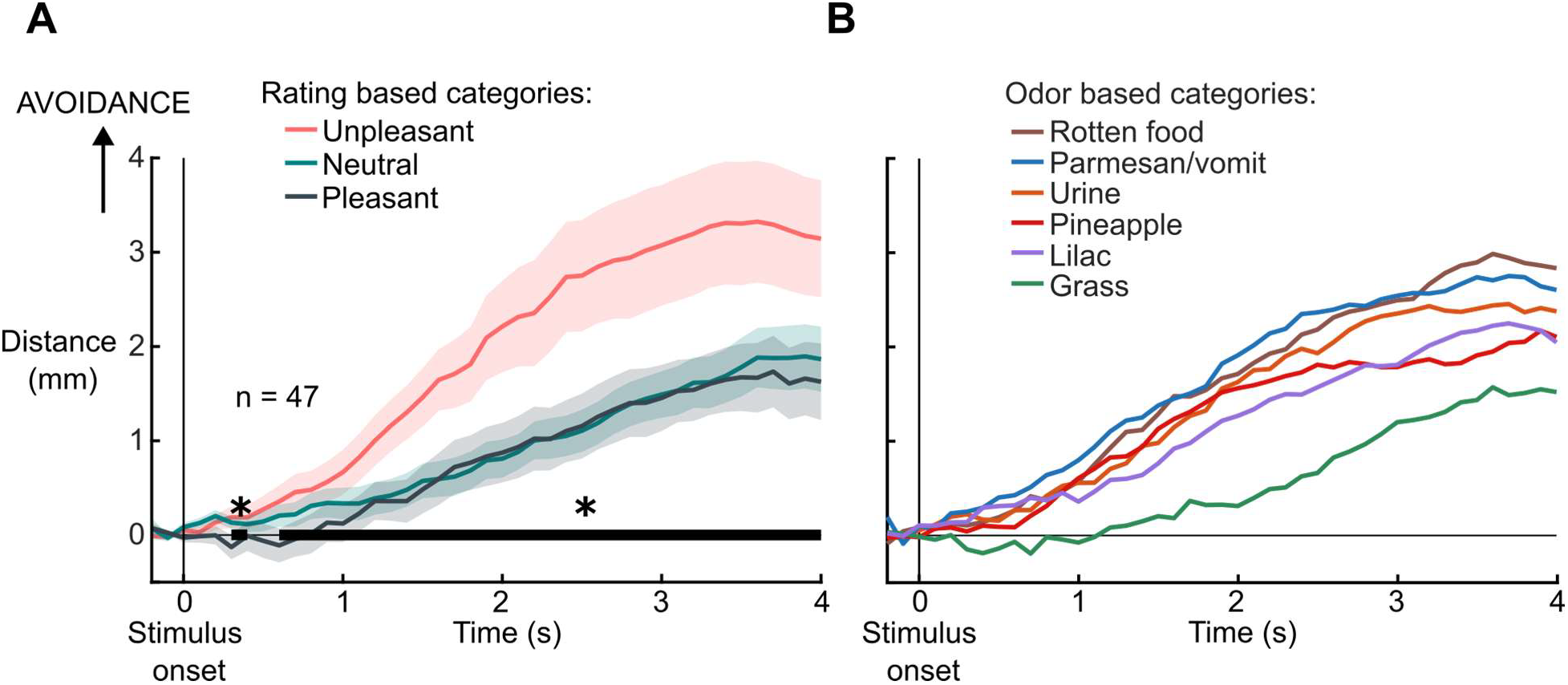
Postural reactions to odors. A) Postural reactions to Unpleasant, Neutral, and Pleasant odors as defined by participants’ individual trial-by-trial valence ratings. The black thick line marks a cluster of time points of statistically significant differences between Unpleasant and Pleasant trials. Stimulus duration was 2 s. Error bars depict ±1 SEM. B) Postural reactions to the 6 different odors. Note that each odor can be rated differently both across participants and across trials for the same participant and can thus contribute to any or all of the conditions in A.

For comparison, we also grouped the trials by odor identity (Fig. 3B). Unsurprisingly, urine, rotten food, and parmesan/vomit tended to be perceived as less pleasant than grass, pineapple, and lilac (urine: M =3.10, SD = 0.96, rotten food: M = 3.21, SD = 1.44, parmesan/vomit: M = 3.42, SD = 1.04, grass: M = 4.93, SD = 1.28, pineapple: M = 6.01, SD = 1.36, and lilac: M = 6.55, SD = 1.26), and this was quantitatively reflected in the pattern of postural reactions (Figure 3B); the three most unpleasant odors elicited the largest avoidance reactions. The perceived intensities were comparable across the six odors, apart from grass which was perceived as slightly less intense: Urine M = 5.26, SD = 1.66, rotten food: M = 6.27, SD = 1.27, parmesan/vomit: 5.91, SD = 1.45, grass: 4.39, SD = 1.50, pineapple: 6.18, SD = 1.16, and for lilac: M = 6.25, SD = 1.14).

For all three categories (Unpleasant, Neutral, and Pleasant), there is an overall trend of moving away from the camera/odor source (Fig. 3A) potentially reflecting the participants’ movement related to their inhalation (the odors were released during inhalation).

## Discussion

We here introduce an ecologically valid and unobtrusive measure of approach–avoidance behavior. Using 3D tracking, we demonstrate postural avoidance reactions to subjectively unpleasant stimuli in both the visual and olfactory domains. This universal avoidance response across sensory modalities confirms the validity and general applicability of our novel measure. Consistent with previous work showing that avoidance reactions tend to be stronger and more reliable than approach reactions, we found pronounced avoidance reactions to unpleasant stimuli as compared to pleasant and neutral stimuli and only minor approach reactions to pleasant stimuli. This asymmetry is possibly reflecting the greater adaptive value of responding rapidly to potential imminent threats compared to pursuing rewards; an interpretation that is consistent with electrophysiological findings showing that unpleasant odors have a direct and privileged access to the olfactory bulb that is not observed for pleasant odors (Iravani et al., 2021; Nordén et al., 2024).

A key strength of the demonstrated 3D camera-based method is that it allows monitoring of natural reactions while keeping participants naïve to the purpose of the experiment, overcoming the limits of traditional methods. The latter either direct participants’ attention to their posture or involve cognitive tasks; for example, participants are asked to perform a task directly linked to approach-avoidance or to stand on a force-plate, making it difficult to hide the aim of the study. These common measures of approach avoidance reactions risk reflecting conscious strategies, demand characteristics, or task compliance. In contrast, the presented 3D camera setup allows participants to behave naturally, and the recorded movements likely reflect non-conscious motor output of underlying motivational processes. We argue that our measurement of distance to the nose is a direct and ecologically valid measure of approach–avoidance behavior. Unlike force-plate measures, which primarily index shifts in whole-body posture and may miss avoidance responses that preserve overall balance (e.g., turning the head away), our method captures a genuine act of avoiding the stimulus. Specifically, it measures movement of the nose toward or away from the source, which more directly aligns with how organisms naturally sample or avoid nearby stimuli.

A limitation of our method is that even though postural displacement provides a sensitive index of avoidance behavior, it does not capture other parts of the defensive response, such as e.g. changes in muscle tone or autonomic adjustments. Additionally, although our results generalize across two sensory modalities, future studies should examine whether similar patterns emerge for other forms of affective stimulation and in more dynamic or socially interactive contexts. The occasional failure of the face-detection algorithm was primarily attributable to lighting conditions and reflections; future work should improve these conditions of the room to optimize the results of the technique. The previously demonstrated effect of deeper inhalation in response to pleasant odors (e.g. (Bensafi et al., 2007) does not constitute a potential confound in our olfactory experiment data, as it would counteract our observed results of larger avoidance for Unpleasant stimulation. This is because we observed an overall effect of moving away from the odor source in response to stimulation of all valence levels, Unpleasant, Neutral, and Pleasant (and to all six odors), indicating that it likely reflects a body movement caused by the inhalation to which odor delivery was synchronized, but where unpleasant odor triggered an avoidance response beyond this inhalation-related effect.

The temporal dynamics of our observed avoidance responses support the notion that unpleasant stimuli trigger an automatic reaction to move the individual away from the source. In both studied sensory modalities, significant differences between unpleasant and pleasant conditions emerged within a time window compatible with early perceptual processing and the initiation of motor adjustments. For olfaction, the delay (at 300 ms, and a continuous effect with onset at 600 ms) is consistent with known delays in odor transduction and cortical processing (∼400 ms following stimulus onset) (Bae et al., 2021; Iravani et al., 2020; Mainland et al., 2014) and aligns with past results using a force-place setup (Iravani et al., 2021). We argue that our results reflect olfactory valence information being swiftly translated into motor biasing signals that promote withdrawal from potentially harmful stimuli. Previous studies have shown that unpleasant odors can elicit fast avoidance-related responses such as changes in sniffing behavior, facial muscle activations, and motor action within a similar time frame (Arshamian et al., 2017; Bensafi et al., 2007; Iravani et al., 2021). The avoidance response to unpleasant visual stimuli emerged later, at 1.5 s after stimulus onset. While affective visual information is extracted rapidly, the translation of this information into whole-body postural adjustments may require sustained stimulus exposure, particularly in the absence of explicit action demands (Bradley et al., 2001; Perakakis et al., 2012). The difference in the timing of the observed visual and olfactory avoidance responses thus likely reflects differences in ecological relevance, sensorimotor coupling, and motor implementation rather than a fundamentally different underlying mechanism or affective evaluation. Compared to visual input, unpleasant odors more likely represent immediate and proximal threats, such as contamination or toxicity, which require rapid whole-body motor adjustments (Boesveldt & Parma, 2021; Stevenson, 2010). All in all, these dynamics are in line with the idea that emotional evaluation biases whole-body motor behavior even in the absence of explicit task demands.

In the visual experiment, the avoidance effect was less pronounced when we grouped trials using normative valence categories, underscoring that although there is some level of consensus, there is an added sensitivity gained by accounting for subjective experience. Similarly, in the olfactory experiment, the three odors with the lowest average valence ratings (urine, rotten food, and butyric acid) elicited qualitatively larger average avoidance reactions than the remaining three odors with higher average pleasantness ratings (grass, lilacs, and pineapple).

In conclusion, the present findings provide robust evidence for automatic avoidance responses to subjectively unpleasant stimuli and demonstrate the utility of the novel 3D camera-based method for capturing these responses in humans. The consistent avoidance effects across vision and olfaction suggest that they reflect a modality-independent motivational mechanism. By linking individual emotional experience to non-conscious motor behavior across sensory domains, this approach offers a promising tool for advancing the study of emotion-action coupling in ecologically valid settings.

## Acknowledgements

This work was supported by the Knut and Alice Wallenberg Foundation (KAW 2018.0152) and the ERC Synergy grant D2Smell (Digitising Smell: From Natural Statistics of Olfactory Perceptual Space to Digital Transmission of Odors) awarded to JNL, and by a Swedish Research Council Grant (2021-03184) awarded to AT.

## Supplementary material

### IAPS images

The IAPS images used as stimulation in Experiment 1 were based on normative data to form categories Unpleasant, Neutral, and Pleasant.

The 36 unpleasant images from the IAPS database were: 1050, 1114, 1120, 1300, 1302, 1930, 1932, 2345.1, 2703, 2730, 3500, 6212, 6230, 6260, 6312, 6313, 6315, 6350, 6510, 6520, 6540, 6550, 6563, 7359, 7380, 9050, 9075, 9163, 9183, 9185, 9187, 9491, 9560, 9622, 9623, 635. The 36 neutral images from the IAPS database were: 2026, 2036, 2190, 2480, 2520, 2745, 5500, 7000, 7003, 7004, 7006, 7009, 7010, 7012, 7014, 7019, 7020, 7025, 7026, 7035, 7041, 7045, 7050, 7052, 7056, 7077, 7095, 7140, 7150, 7160, 7233, 7235, 7242, 7495, 7710, 8170.

The 36 pleasant images from the IAPS database were images number: 2045, 2058, 2071, 2080, 2150, 2158, 2303, 2344, 2345, 2346, 4597, 4599, 4607, 4608, 4611, 4660, 5260, 5270, 5450, 5480, 5621, 5628, 5700, 5910, 7405, 8030, 8080, 8178, 8179, 8185, 8186, 8210, 8370,8490, 8492, 8496.

